# BMP Signaling Downstream of the Highwire E3 Ligase Sensitizes Nociceptors

**DOI:** 10.1101/251991

**Authors:** Ken Honjo, W. Daniel Tracey

## Abstract

A comprehensive understanding of the molecular machinery important for nociception is essential to improving the treatment of pain. Here, we show that the BMP signaling pathway regulates nociception downstream of the E3 ubiquitin ligase *highwire* (*hiw*). *Hiw* loss of function in nociceptors caused antagonistic and pleiotropic phenotypes with simultaneous insensitivity to noxious heat but sensitized responses to optogenetic activation of nociceptors. Thus, *hiw* functions to both positively and negatively regulate nociceptors. We find that a sensory transduction-independent sensitization pathway was associated with BMP signaling. BMP signaling in nociceptors was up-regulated in *hiw* mutants, and nociceptor-specific expression of *hiw* rescued all nociception phenotypes including the increased BMP signaling. Blocking the transcriptional output of the BMP pathway with dominant negative Mad suppressed nociceptive hypersensitivity that was induced by interfering with hiw. The up-regulated BMP signaling phenotype in *hiw* genetic mutants could not be suppressed by mutation in *wallenda* suggesting that *hiw* regulates BMP in nociceptors via a *wallenda* independent pathway. In a newly established Ca^2+^ imaging preparation, we observed that up-regulated BMP signaling caused a significantly enhanced Ca^2+^ signal in the axon terminals of nociceptors that were stimulated by noxious heat. This response likely accounts for the nociceptive hypersensitivity induced by elevated BMP signaling in nociceptors. Finally, we showed that acute activation of BMP signaling in nociceptors was sufficient to sensitize nociceptive responses to optogenetically-triggered nociceptor activation without altering nociceptor morphology. Overall, this study demonstrates the previously unrevealed roles of the Hiw-BMP pathway in the regulation of nociception and provides the first direct evidence that up-regulated BMP signaling physiologically sensitizes responses of nociceptors and nociception behaviors.

**Author Summary:** Although pain is a universally experienced sensation that has a significant impact on human lives and society, the molecular mechanisms of pain remain poorly understood. Elucidating these mechanisms is particularly important to gaining insight into the clinical development of currently incurable chronic pain diseases. Taking an advantage of the powerful genetic model organism *Drosophila melanogaster* (fruit flies), we unveil the Highwire-BMP signaling pathway as a novel molecular pathway that regulates the sensitivity of nociceptive sensory neurons. Highwire and the molecular components of the BMP signaling pathway are known to be widely conserved among animal phyla, from nematode worms to humans. Since abnormal sensitivity of nociceptive sensory neurons can play a critical role in the development of chronic pain conditions, a deeper understanding of the regulation of nociceptor sensitivity has the potential to advance effective therapeutic strategies to treat difficult pain conditions.

## Introduction

In spite of its clear medical importance, the molecular mechanisms of pain signaling remain poorly understood. Pain pathways in large part depend on sensory input from specialized sensory neurons called nociceptors (1). Since the activation of nociceptors leads to pain sensation and the sensitization of nociceptors is thought to be a major contributor of pain pathogenesis, understanding the molecular mechanisms controlling nociceptor function is essential for improving the treatment of pain (2).

*Drosophila melanogaster* is a powerful model system for neurogenetic studies of nociception. Larval *Drosophila* show stereotyped behavioral responses to potentially tissue-damaging stimuli, such as noxious heat or harsh mechanical stimulation (3). The most unambiguous larval nociception behavior involves a corkscrew like rolling around the long body axis (termed nocifensive escape locomotion (NEL) or simply “rolling”). Since rolling is specifically triggered by noxious stimuli and is clearly separable from normal larval locomotion, the analysis of NEL provides a robust behavioral paradigm to study nociception. Class IV multidendritic (md) neurons are polymodal nociceptors that are necessary for thermal and mechanical nociception in larvae (4). Optogenetic activation of the Class IV neurons is sufficient for triggering NEL(4, 5). Accumulating evidence in studies of fly nociception suggests that the molecular pathways of nociception are conserved between *Drosophila* and mammals (3, 6-15).

To identify genes important for nociceptor function, we recently performed thermal nociception screens in which we targeted the RNAi knockdown of nociceptor-enriched genes in a nociceptor-specific manner (16). In this screen, we found that two RNAi lines targeting *highwire* (*hiw*) caused driver dependent hypersensitivity in thermal nociception assays (revealed as a rapid response to a threshold heat stimulus) indicating a potential role for *hiw* as a negative regulator of nociceptor activity (16). *hiw* is an evolutionally conserved gene encoding an E3 ubiquitin ligase, whose function has been implicated in various aspects of neuronal development, synaptic function, and neuronal degeneration (17). However, in contrast, very little is known about the roles of *hiw* in sensory processing and in controlling behavior. Here, we present additional and more specific evidence that *hiw* plays an important role in the regulation of behavioral nociception and nociceptor sensitivity through the bone morphogenetic protein (BMP) pathway.

## Results

### Highwire regulates the sensitivity of nociceptors

To further investigate the potential function of *hiw* in nociception that was suggested by our previous study, we tested mutants for a strong loss-of-function allele of *hiw* (*hiw^ND8^*) in thermal nociception assays (18). Unexpectedly, we found that genetic mutants of *hiw* showed insensitivity to a noxious temperature probe of 42 or 46°C, which was, surprisingly, the opposite of the previously described *hiw* RNAi phenotype (Fig 1A and data not shown). Similar thermal insensitivity was also seen with other *hiw* alleles (Fig S1). Although *hiw* is widely expressed in the nervous system (18), nociceptor-specific restoration of *hiw* expression rescued this insensitivity (Fig 1A), indicating that *hiw* function in nociceptors is sufficient for restoration of normal thermal nociception and the relevant site of action was in nociceptors.

**Fig 1.**
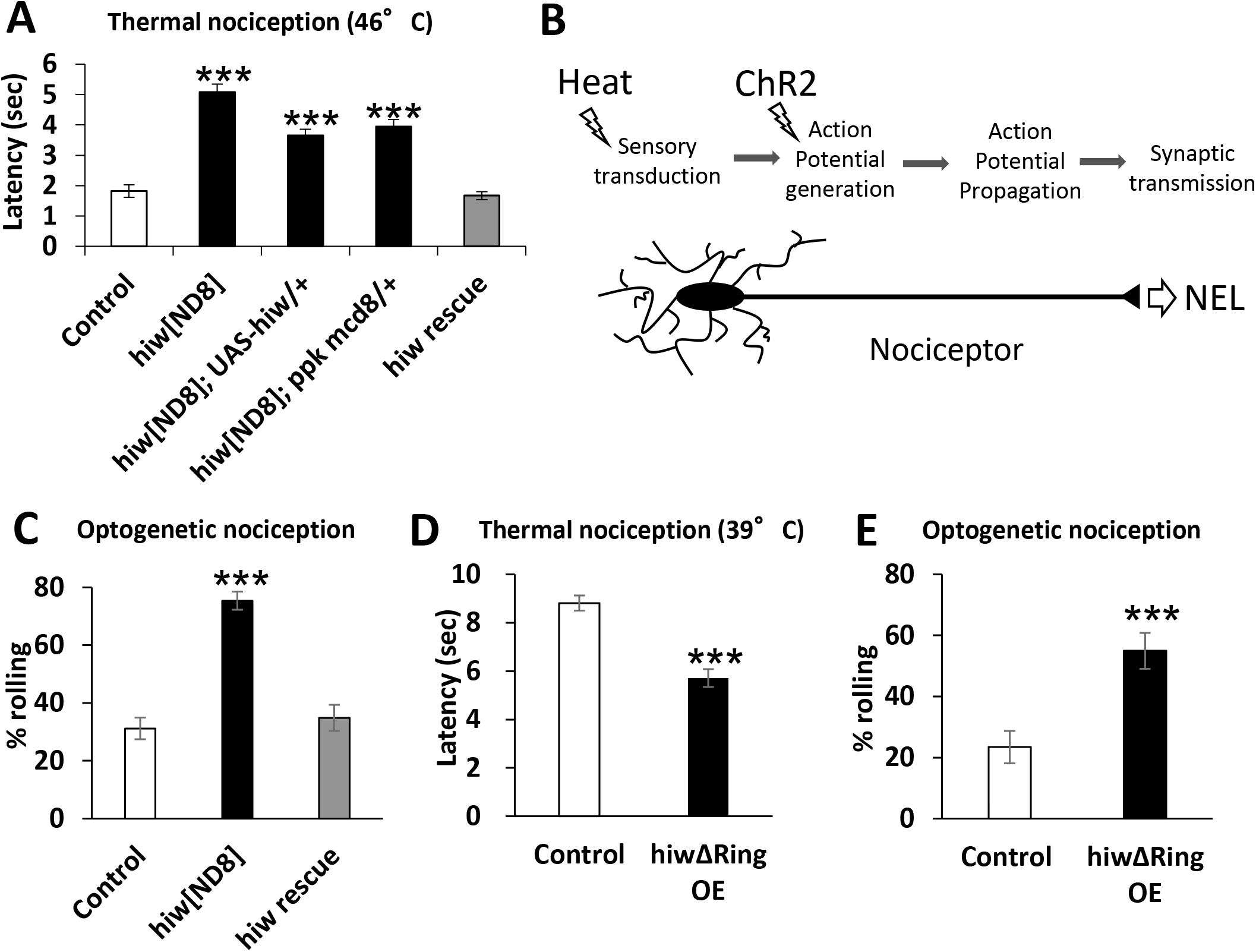
*hiw* is involved in both desensitizing and sensitizing pathways in nociceptors. (A) Insensitive thermal nociception in *hiw^ND8^* mutant and nociceptor-specific rescue of the insensitivity. In comparison to the control *w^1118^* (n = 119, 1.8 ± 0.2), *hiw^ND8^*(n = 114, 5.1 ± 0.3), no driver control (*hiw^ND8^*; UAS-*hiw*/+, n = 108, 3.7 ± 0.2) and *hiw^ND8^* with GAL4 driver (*hiw^ND8^*; *ppk*-*GAL4 UAS*-*mCD8::GFP*/+, n = 101, 3.9 ± 0.2) all showed significantly delayed nociceptive responses to a 46°C probe, while the rescue genotype (*hiw^ND8^*; *ppk-GAL4 UAS-mCD8::GFP*/*UAS-hiw*, n = 122, 1.7 ± 0.1) had a normal response. *** p < 0.001 (Steel's test versus control). (B) A schematic of thermal and optogenetic stimulation of a nociceptor. While heat stimuli activate nociceptors via sensory transduction, ChR2 triggers nociceptor activation independently of sensory transduction. (C) The *hiw* genetic mutant expressing ChR2 in nociceptors was more responsive than the control to optogenetically triggered nociceptor activation, and the nociceptor-specific expression of *hiw* rescued optogenetic nociception responses to levels similar to control. Control (*w^1118^*/Y; *ppk*-*GAL4 UAS*-*ChR2::YFP*/+, n = 154, 31 ± 4%), *hiw^ND8^* (*hiw^ND8^*/*Y*; *ppk*-*GAL4 UAS-ChR2::YFP*/+, n =191, 75 ± 3%) and *hiw* rescue (*hiw^ND8^*/*Y*; *ppk*-*GAL4 UAS-ChR2::YFP*/*UAS-hiw*, n = 112, 35 ± 5%). *** p < 0.001 (Fisher's exact test with Bonferroni correction). (D) *hiwΔRing* expression (*hiwΔRing* OE) in nociceptors resulted in thermal hypersensitivity. *hiwΔRing* OE animals (*ppk-GAL4* x *UAS-hiwΔRing*, n = 90, 5.7 ± 0.4) showed a significantly shortened latency to respond to a 39°C thermal probe compared to controls (ppk-GAL4 x *w^1118^*, n = 104, 8.8 ± 0.3). *** p < 0.001 (Mann-Whitney's U-test). (E) *hiwΔRing* expression in nociceptors induced hypersensitivity to optogenetic nociceptor stimulation. *hiwΔRing*-expressing animals (*ppk*-*GAL4 UAS-ChR2::YFP x UAS*-*hiwΔRing*, n = 71, 55 ± 6%) exhibited increased responsiveness to optogenetic stimulation of nociceptors compared to control animals (*ppk*-*GAL4 UAS-ChR2::YFP x w^1118^*, n = 64, 23 ± 5%). *** p < 0.001 (Fisher's exact test). All error bars represent standard error.

Intrigued by the clear phenotypic distinction between genetic mutants and RNAi animals, we further dissected the nociception phenotype of *hiw* mutants by employing an optogenetic strategy. Optical activation of larval nociceptors via the blue light-gated cation channel Channelrhodopsin-2 (ChR2) is sufficient to induce larval NEL (4, 5). Since nociceptor activation by ChR2 circumvents sensory transduction but still depends on the machinery essential for downstream signaling (Fig 1B), this technique has been utilized to distinguish genes that are important for primary sensory function from those that function in downstream aspects of signaling, such as action potential generation/propagation and/or synaptic transmission (10, 19). Using low intensity blue light (3.8 klux), which elicits NEL in about 20-30% of control animals expressing ChR2::YFP in nociceptors (Fig 1C), we found that the *hiw^ND8^* mutants had a significantly increased probability to show NEL, indicating that the mutant for this allele is hypersensitive in response to optogenetic activation of nociceptors (Fig 1C) even though it was insensitive in thermal nociception assays. Tissue specific rescue experiments again showed that nociceptor specific expression of *hiw* was sufficient to rescue this optogenetic hypersensitivity (Fig 1C). Taken together, these findings suggested that *hiw* has multiple, but dissociable, effects in the regulation of nociceptors. On the one hand, *hiw* regulated a sensory transduction-dependent function causing insensitivity, but it also regulated a function further downstream of sensory transduction that caused hypersensitivity. Thus, the hypersensitivity seen in our earlier RNAi experiments is likely reflective of effects on the latter process.

To further examine hiw's role, we tested the effects of expressing *hiwΔRING* in nociceptors. The *hiwΔRING* transcript encodes a mutated form of *hiw* lacking the RING domain that is responsible for E3 ligase activity (20, 21). This mutated protein has been proposed to function as a dominant-negative poison subunit in multimeric Hiw E3 ligase complexes. Similar to our original observations with *hiw RNAi*, expression of *hiwΔRING* in nociceptors resulted in significant hypersensitivity in thermal nociception (Fig 1D and S2). This manipulation also caused hypersensitive optogenetic nociception responses (Fig 1E). As *hiw* encodes a large protein with many functional domains, and phenotypes of *hiw* mutants are known to show varied sensitivity to gene dosage (Wu et al., 2005), the observed similarity between *hiwΔRING* overexpression and *hiw* RNAi is suggestive of dosage-dependent effects of *hiw* in nociceptors. For instance, the dominant negative approach may lead to an incomplete loss of function for *hiw* that is similar to the effects of *RNAi*.

### Hiw attenuates BMP signaling in nociceptors

It has been very recently shown that the canonical BMP pathway in nociceptors is required for nociceptive sensitization after tissue damage in *Drosophila* (22). Since the BMP signaling pathway has also been proposed to be a downstream pathway regulated by Hiw in motoneurons (23), we tested whether the BMP signaling pathway is regulated downstream of Hiw in nociceptors. We first examined the level of phosphorylated Mad (pMad) in nociceptor nuclei by quantitative immunohistochemistry, which is an established method for evaluating the activation level of intracellular BMP signaling (24-30). In nociceptor nuclei, *hiw* genetic mutants showed significantly elevated pMad levels (33%) in comparison to wild-type, even when processed together in the same staining solution (see also Materials and Methods) (Fig 2 A, B and F). A similarly modest change in pMad accumulation in motor neuron nuclei is associated with effects on presynaptic function and morphology at the neuromuscular junction (NMJ) (31, 32). An increased accumulation of pMad in the nucleus and the cytoplasm was observed in nociceptors expressing *hiwΔRING* (Fig 2C and F). Expression of wild-type *hiw* in nociceptors of *hiw* mutant animals rescued the elevated pMad level (Fig 2D and F). We also confirmed that our immunohistochemistry successfully detected the increase of nuclear pMad caused by expressing the constitutively active form of *thick veins* (*tkv^QD^*), which activates the intracellular BMP signaling cascade independently of BMP ligands (33) (Fig 2E and F). These data together suggest that BMP signaling is negatively regulated downstream of *hiw* in larval nociceptors. In the larval motoneurons, it is known that pMad signals can be locally detected at synaptic boutons as well as nuclei (25, 34, 35). However, no detectable pMad signals were observed at synaptic terminals in larval nociceptors (Fig 2G).

**Fig 2.**
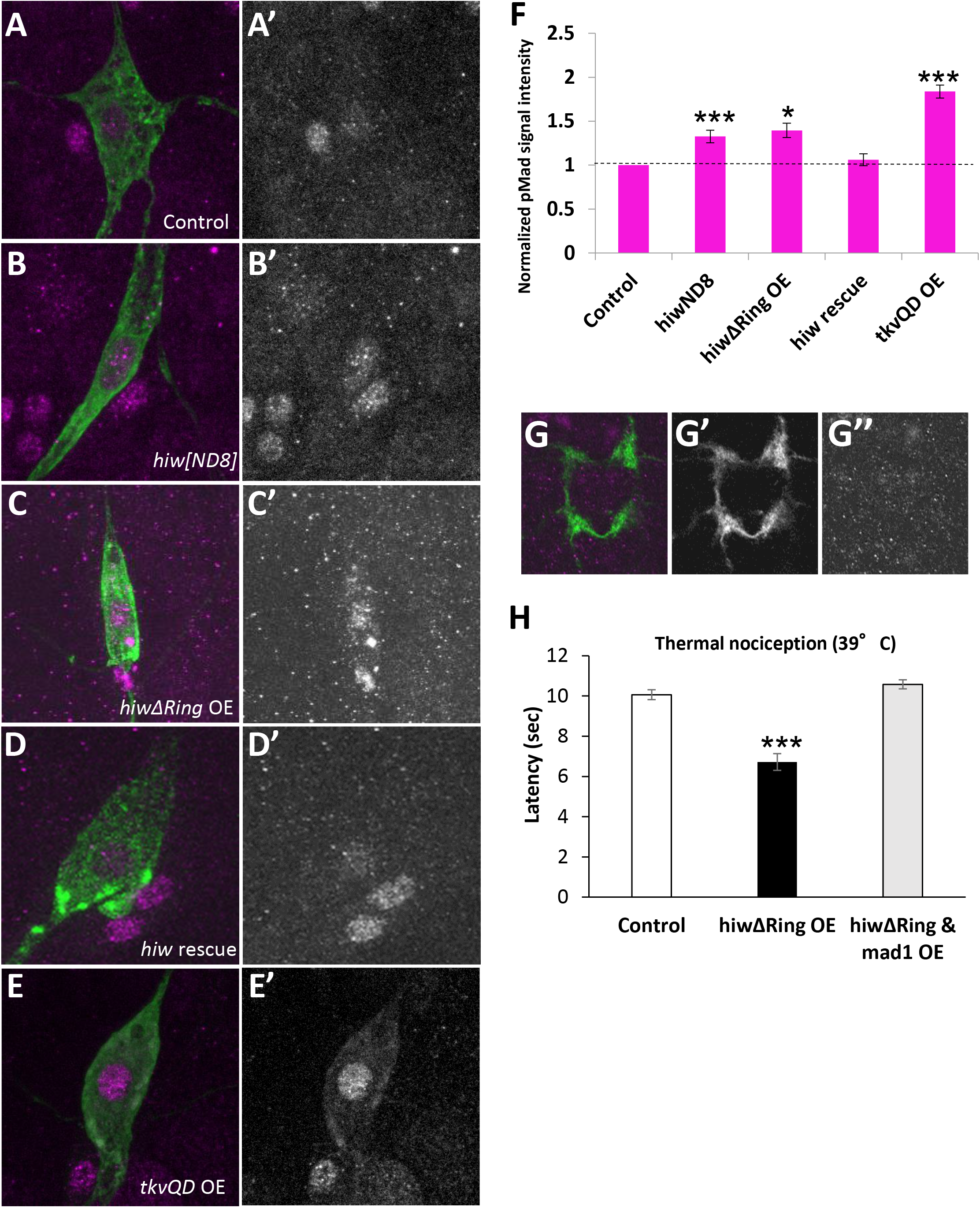
BMP signaling in nociceptors is negatively regulated at the downstream of *hiw*. (A-E) Representative images of pMad immunoreactivity in nociceptor cell bodies. Green represents mCD8::GFP and magenta shows pMad signals. (A’-E’) Split images for pMad signals. (F) Quantification of nuclear pMad signals in nociceptors. *hiw^ND8^* mutants (*hiw^ND8^*; *ppk-GAL4 UAS-mCD8::GFP*/+, n = 21) and *hiwΔRing* OE (*ppk-GAL4 UAS-mCD8::GFP*/+; *UAS-hiwΔRing*/+, n = 18) had 33 ± 7% and 40 ± 8% increases in nuclear pMad signals, respectively. No significant difference in nuclear pMad level compared to controls was detected in *hiw rescue* animals (*hiw^ND8^*; *ppk-GAL4 UAS*-*mCD8::GFP*/*UAS-hiw*, n = 24). Nociceptors expressing *tkv*^QD^ (*ppk-GAL4 UAS*-*mCD8::GFP x UAS*-*tkv*^Q*D*^, n = 24) also showed significantly increased nuclear pMad levels (84 ± 7%). Control (*ppk-GAL4 UAS-mCD8::GFP*/+, n > 24) * p < 0.05, *** p < 0.001 (Mann-Whitney's U-test). (G) A projection image of axon terminal of Class IV neurons at A4 and A5 segments in the larval ventral ganglia. Green represents mCD8::GFP and magenta shows pMad signals. pMad signals at axon termini in nociceptors were not distinguishable from the background. (G’) Split image for mCD8::GFP. (G’’) Split image for pMad signals. (H) Expression of mad^1^ suppressed the thermal hypersensitivity in *hiw4Ring*-expressing animals. Control (*w^1118^ x ppk-GAL4*, n = 73, 10.1 ± 0.3), *hiwΔRing* OE (*ppk-GAL4 x UAS*-*hiwΔRing*, n = 59, 6.7 ± 0.4) and *hiwΔRing* & *mad1* OE (*ppk-GAL4 UAS*-*hiwΔRing x UAS*-*mad*^1^, n = 54, 10.6 ± 0.2). *** p < 0.001 (Steel's test versus control). Error bars represent standard error.

Next we tested whether up-regulated BMP signaling in nociceptors is responsible for the hypersensitive nociceptive responses caused by *hiw* loss-of-function. *mad^1^* encodes a dominant-negative form of Mad with disrupted DNA-binding ability (36). When *mad^1^* was expressed together with *hiwΔRING* in nociceptors, the hypersensitive phenotype that was normally induced by the expression of *hiwΔRING* alone was not detected (Fig 2H). Since neither expressing Mad^1^ together with *hiwΔRING* nor expressing Mad^1^ alone in nociceptors induced insensitivity to noxious heat (Fig S3), these results indicate that hypersensitive nociception caused by weak *hiw* loss of function requires an intact BMP signaling pathway that normally operates through Mad. This result is consistent with the elevated pMad observed with *hiw* loss of function as playing a causal role in the hypersensitive phenotypes.

The MAP kinase kinase kinase (MAPKKK) *wallenda* (*wnd*) is a well-characterized target substrate of Hiw ligase (17). Hiw negatively regulates the protein level of Wnd, and the Hiw-Wnd interaction is crucial for normal synaptic growth, but not for normal synaptic function in NMJ (30, 37-39). In addition, *hiw* interacts with *wnd* in Class IV neurons in the regulation of dendritic and axonal morphology (40). In larval motoneurons, it has been suggested that *wnd* is not involved in the regulation of BMP signaling (30). To test whether *wnd* is involved in the control of BMP signaling downstream of *hiw* in nociceptors, we examined a genetic interaction between *hiw* and *wnd* in double mutants. A *wnd* mutation in *hiw* mutant background did not suppress the elevated nuclear pMad level in nociceptors that we observed in the *hiw* mutant (Fig 3A-D and F), nor did *wnd* single mutants show altered nuclear pMad accumulation relative to controls (Fig 3E and F). Interestingly, significant up-regulation of nuclear pMad signal was observed in nociceptors overexpressing *wnd*, but not with a kinase-dead version of *wnd* (Fig S4). Taken together, these results suggest that elevated nuclear pMad in *hiw* mutant nociceptors does not depend on the activity of Wnd, although overexpression of *wnd* with GAL4/UAS can cause elevated BMP signaling in nociceptors.

**Fig 3.**
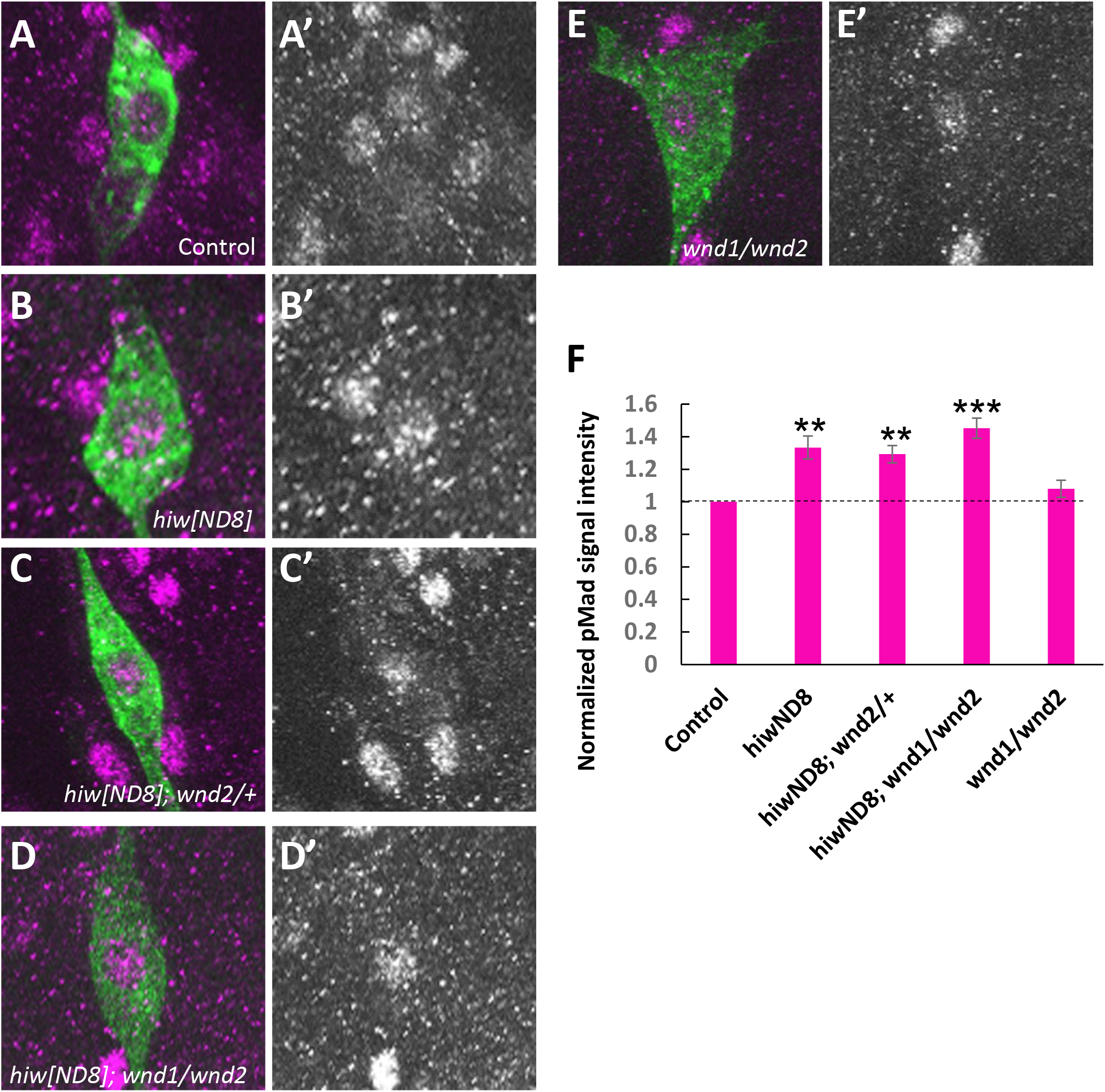
Activated BMP signaling in *hiw* mutant does not depend on *wallenda*. (A-E) Representative images of pMad immunoreactivity in nociceptor soma. Green shows mCD8::GFP and magenta represents pMad signals. (A’-E’) Split images for pMad signals. (F) Quantification of nuclear pMad signals in nociceptors. Similarly to *hiw^ND8^* mutants (*hiw^ND8^*; *ppk*-*GAL4 UAS*-*mCD8::GFP*/+, n = 36), *hiw^ND8^* with heterozygous or transheterozygous *wnd* mutations (*hiw^ND8^*; *ppk*-*GAL4 UAS*-*mCD8::GFP*/+; *wnd^2^*/+ and *hiw^ND8^*; *ppk*-*GAL4 UAS*-*mCD8::GFP*/+; *wnd^1^*/*wnd^2^*, n = 48 and 45) showed significantly increased nuclear pMad level relative to controls (*ppk-GAL4 UAS-mCD8::GFP*/+, n = 48). The transheterozygous *wnd* mutants (*ppk-GAL4 UAS-mCD8::GFP*/+; *wnd^1^*/*wnd^2^*, n = 33) did not show a significant difference in nuclear pMad level compared to controls (p > 0.7). ** p < 0.01, *** p < 0.001 (Mann-Whitney's U-test). Error bars represent standard error.

To gain insight into which regions of Hiw protein are involved in attenuating BMP signaling in nociceptors, we performed an expression study of a series of Hiw dominant negatives with various deletions, as established by Tian et al. (38) (Fig 4A). Expressing HiwNT (N-terminal half of Hiw) caused a greater than 200% increase in nuclear pMad signals compared to controls (Fig 4B, C and H). HiwCT (C-terminal half of Hiw) and HiwΔRCC1 resulted in 99% and 68% increases in nuclear pMad signals, respectively (Fig 4D, E and H). HiwCT and HiwΔRCC1 also caused marked accumulation of pMad signals in the cytoplasm of nociceptors (Fig 4D and E), which was also observed with HiwΔRing expression (Fig 2C). This cytoplasmic accumulation of pMad signals is unlikely due to technical variability of immunostaining since the control samples treated in the same staining solutions with HiwCT or HiwΔRCC1 never developed such accumulations and cells nearby the nociceptors showed the normal pMad signal. In contrast, HiwΔHindIII and HiwCT1000 (C-terminal only region of Hiw) did not cause any changes in nuclear pMad signals in nociceptors (Fig 4C, F and H). Thus, the attenuation of BMP signaling in nociceptors through Hiw appears to depend on different regions of Hiw from those that have been proposed to be involved in the regulation of NMJ morphology (HiwΔRCC1, and HiwΔHindIII function as dominant-negative in NMJ morphology while HiwNT and HiwCT1000 do not (38)). Because both HiwNT and HiwCT, which are largely non-overlapping N-terminal and C-terminal halves of Hiw, caused increased nuclear pMad signals, multiple regions of the Hiw protein must be intact for normal suppression of BMP signaling in nociceptors.

**Fig 4.**
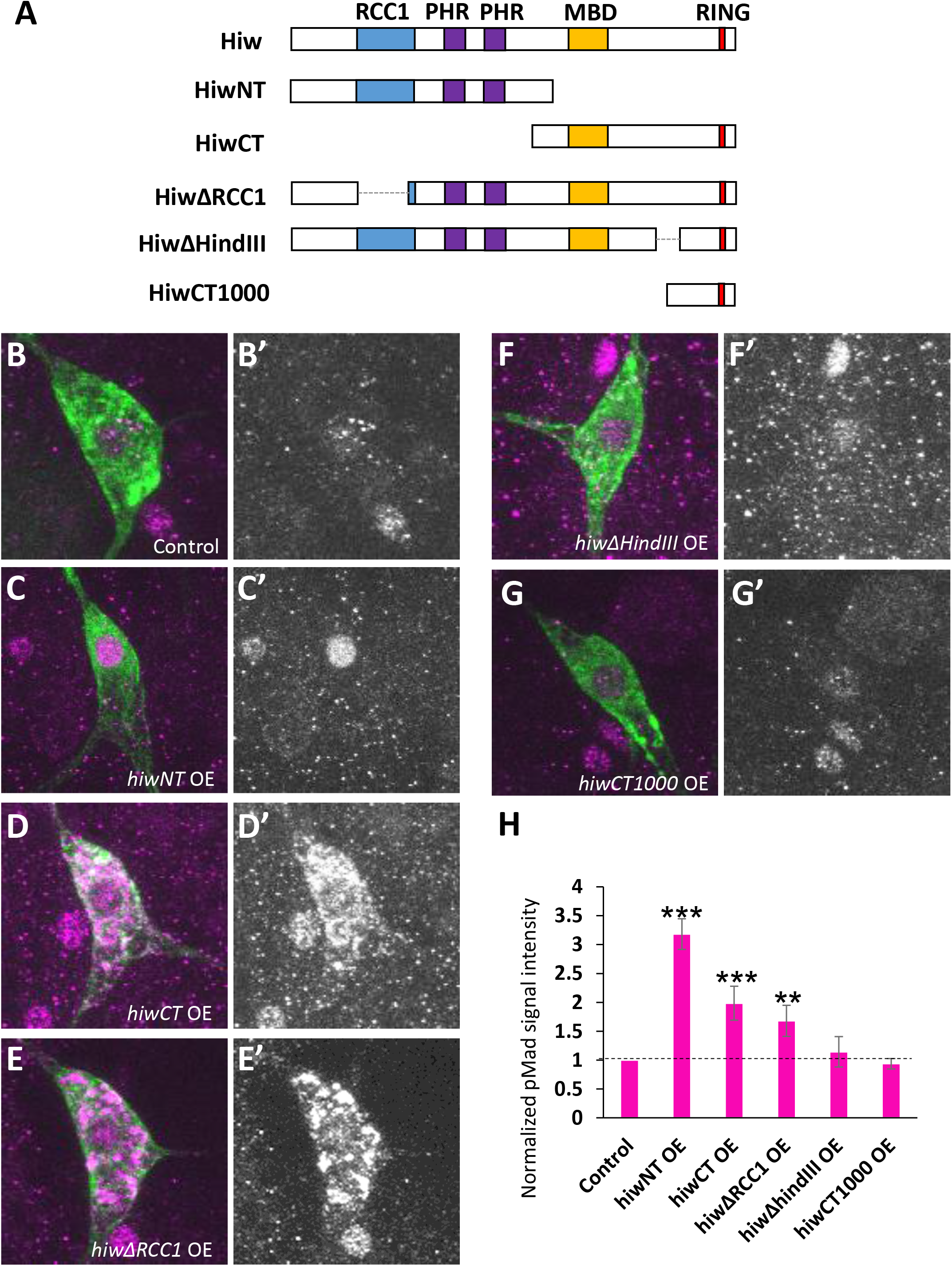
pMad signals in nociceptors expressing various *hiw* deletion constructs. (A) A schematic showing the structure of Hiw and Hiw deletion constructs. (B-G) Representative images of pMad immunoreactivity in nociceptor cell bodies. Green represents mCD8::GFP and magenta shows pMad signals. (B’-G’) Split images for pMad signals. (H) Quantification of nuclear pMad signals in nociceptors. Nociceptors expressing *hiwNT* OE (*ppk*-*GAL4 UAS*-*mCD8::GFP* x *UAS-hiwNT*, n = 12), *hiwCT* OE *(ppk-GAL4 UAS-mCD8::GFP* x *UAS-hiwCT*, n = 12) and *hiwΔRCC1* OE (*ppk-GAL4 UAS-mCD8::GFP* x *UAS-hiwΔRCC1*, n = 12) showed nuclear pMad signals increased by 218 ± 26%, 99 ± 19% and 68 ± 18%, respectively. A significant difference in nuclear pMad level compared to controls was not detected in *hiwΔHindIII* OE (*ppk-GAL4 UAS-mCD8::GFP* x *UAS-hiwΔHindIII*, n = 12) or *hiwCT1000* OE (*ppk-GAL4 UAS-mCD8::GFP* x *UAS-hiwCT1000*, n = 12). Control (*ppk-GAL4 UAS-mCD8*:: *GFP*/+, n = 12) ** p < 0.01, *** p < 0.001 (Mann-Whitney's U-test). Error bars represent standard error.

### Elevated BMP signaling in nociceptors induces behavioral nociceptive hypersensitivity

Although a previous study by Follansbee et al. suggests that the canonical BMP signaling pathway in larval nociceptors is a necessary component for nociceptive sensitization after tissue-damage, whether up-regulation of BMP signaling in nociceptors is sufficient to sensitize nociception has not been proven and potential mechanisms leading to sensitization are unknown. Because our data support the notion that the up-regulation of BMP signaling in nociceptors plays a key role in inducing sensitized nociception, we tested whether up-regulation of intracellular BMP signaling in nociceptors is sufficient to induce nociceptive hypersensitivity. In thermal nociception assays, animals expressing the constitutively active BMP receptor *tkv^QD^* in nociceptors did exhibit significant hypersensitivity (Fig 5A and S2), and *tkv^QD^* also caused hypersensitive responses in optogenetic nociception assays. The latter suggests that elevated BMP signaling in nociceptors was able to sensitize nociception through a mechanism that was downstream of sensory transduction (Fig 5B). Although the dendritic structure of nociceptors in *tkv^QD^* overexpressors was not significantly altered (Fig 5C-E), overexpression of *tkv^QD^* caused overextension and overexpansion of nociceptor axon termini (Fig 5F-H). Combined, these data demonstrate that elevated BMP signaling in nociceptors is sufficient to sensitize thermal and optogenetic nociception behaviors in addition to causing increases in axon terminal branching.

**Fig 5.**
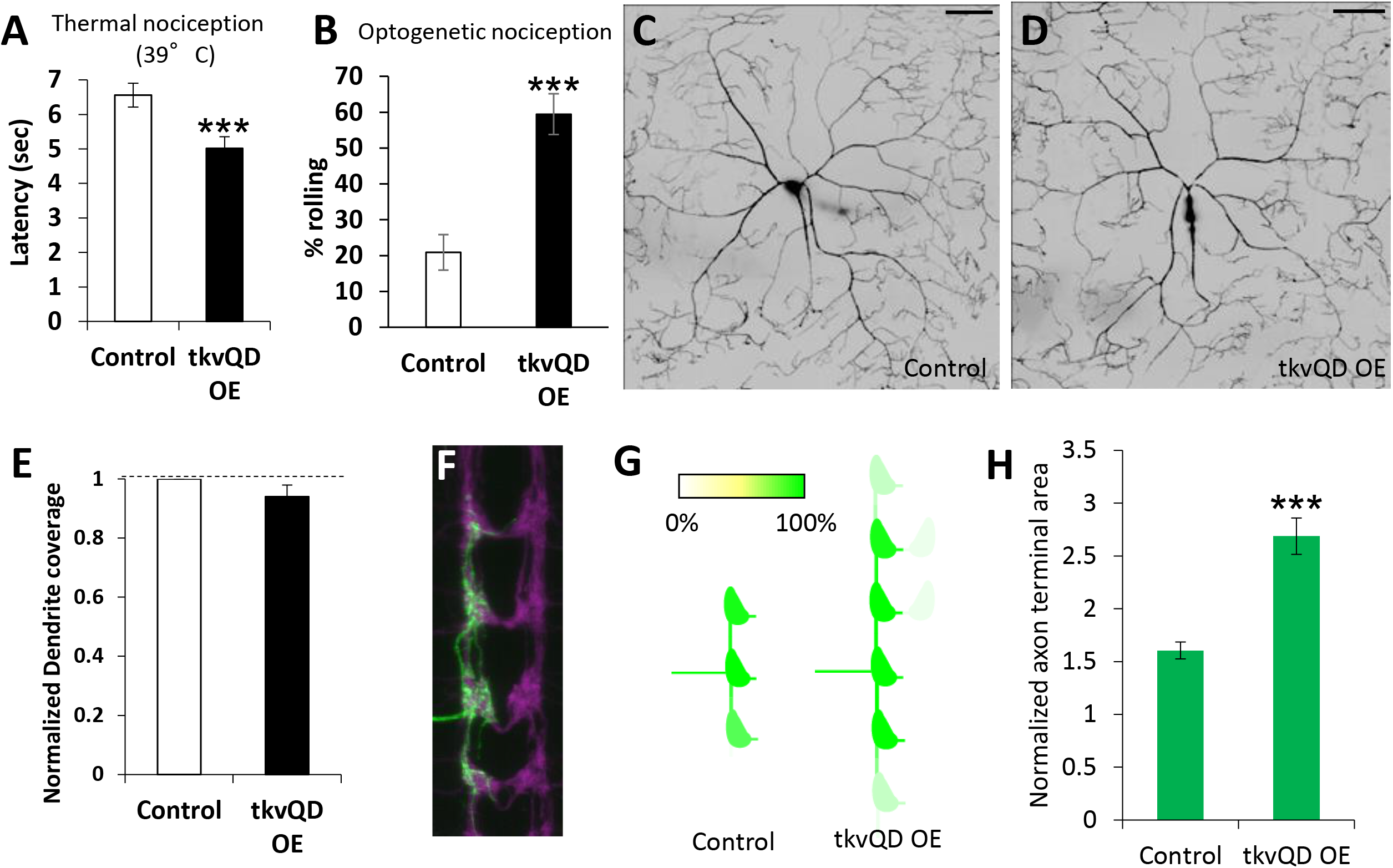
Activation of BMP signaling in nociceptors induces nociceptive hypersensitivity. (A) Animals expressing *tkv*^Q*D*^ in Class IV neurons showed thermal hypersensitivity. Control (*ppk-GAL4* x *w^1118^*, n=102, 6.6 ± 0.3) and *tkv*^Q*D*^ OE (*ppk-GAL4 x UAS-tkv*^QD^, n=118, 5.0 ± 0.3). ** p < 0.01 (Mann-Whitney's U-test). (B) Expression of *tkv*^QD^ in Class IV neurons caused optogenetic hypersensitivity. *tkv*^QD^ overexpressors expressing ChR2::YFP in nociceptors (*ppk-GAL4 UAS-ChR2::YFP x UAS-tkv*^QD^, n = 74, 59 ± 6%) showed significantly elevated responsiveness to blue light-triggered nociceptor activation compared to controls (*ppk-GAL4 UAS-ChR2::YFP*x *w^1118^*, n = 67, 21 ± 5%). *** p < 0.001 (Fisher's exact test). (C-E) Overexpression of *tkv*^QD^ in nociceptors did not affect dendritic coverage. (C and D) Representative images of ddaC dendrites in control (*ppk-GAL4 UAS-mCD8::GFP* x *w^1118^*) and *tkv*^QD^ overexpression (*ppk-GAL4 UAS-mCD8*:: *GFP x UAS-tkv*^QD^) animals. Scale bars represent 100 μm (E) Quantification of dendritic coverage. Dendritic coverage in *tkv*^QD^-overexpressing animals was indistinguishable from that in controls (n = 6 and 6, p > 0.3, Mann-Whitney's U-test). (F-H) Expression of *tkv*^Q*D*^ in nociceptors resulted in overextension of axon termini. (F) A representative image of a v’ada Class IV axon terminal in a *tkv*^Q*D*^ overexpressor (ppk1.9-GAL4; UAS>CD2 stop>mCD8::GFP hs-flp x UAS*-tkv*^Q*D*^). Scale bar represents 5 μm. (G) Heat map of axonal projections. Animals with expression of *tkv*^Q*D*^ showed a severe overextension phenotype (n = 13) compared to controls (ppk1.9-GAL4; UAS>CD2 stop>mCD8::GFP hs-flp x *w^1118^*, n = 24). (H) Quantification of terminal size of the v’ada Class IV neuron. Terminal size of the v’ada axon was significantly increased in *tkv*^Q*D*^-expressing animals (n = 13) compared to controls (n = 24). *** p < 0.001 (Steel's test versus control). All error bars represent standard error.

### Elevated BMP signaling increases Ca^2+^ responses in nociceptor terminals

Since nociceptor-specific up-regulation of BMP signaling sensitizes thermal and optogenetic nociception behaviors, we next explored whether the up-regulation of intracellular BMP signaling actually sensitizes physiological responses of nociceptors. To observe neuronal responses of larval nociceptors to a range of thermal stimuli, we developed a preparation for optical recording from axon terminals of the nociceptive neurons. We then observed these terminals while we locally applied a thermal ramp stimulus to the larval body wall (Fig 6A). To monitor Ca^2+^, the genetically encoded sensor GCaMP6m was expressed under the control of *ppk*-*GAL4* (41). In control animals we observed a steep increase of the GCaMP6m signal in nociceptors when the ramping temperature reached the 39-47°C temperature range (Fig 6B and D). We found that nociceptors expressing *tkv^QD^* showed a significantly greater increase of GCaMP6m signals through 36-50°C in comparison to those in controls (Fig 6C and D), while basal fluorescence levels of GCaMP6m (F_0_) were comparable between the control and *tkv^QD^*-expressing nociceptors (Fig 6E). These results suggest that the significantly greater increase of GCaMP6m signals observed in nociceptors expressing *tkv^QD^* is due to the greater level of Ca^2+^ influx triggered by the heat ramp stimulus, and not to unintended transcriptional upregulation of GCaMP6m. Thus, elevated BMP signaling in nociceptors results in exaggerated Ca^2+^ signals at the terminals of nociceptors in response to heat in the noxious range. This conclusion is consistent with the behavioral nociceptive sensitization induced by the same intracellular up-regulation of BMP signaling in nociceptors.

**Fig 6.**
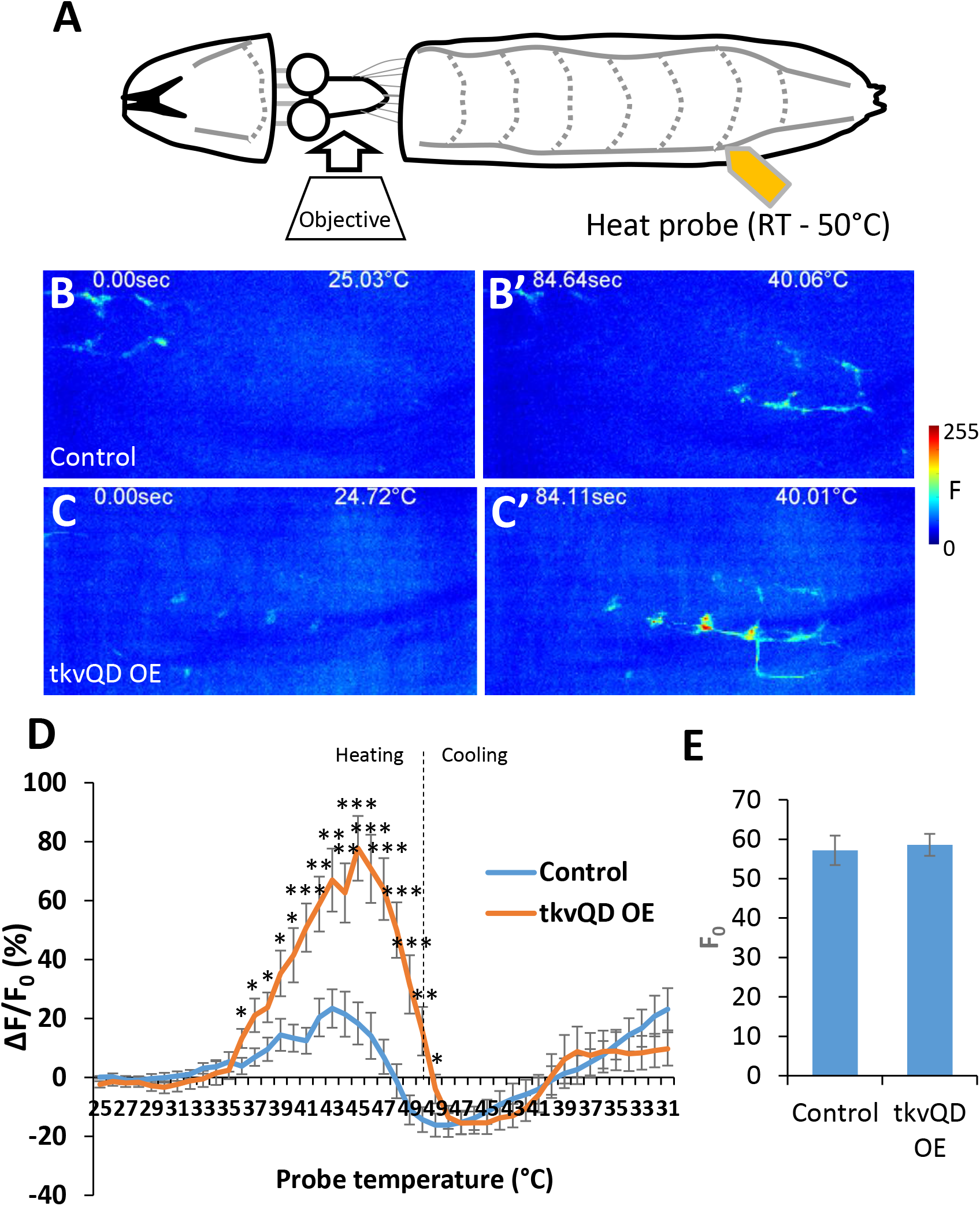
Elevated BMP signaling increases Ca^2^+ responses in nociceptor axon terminals. (A) A cartoon showing the Ca^2+^ imaging preparation to monitor GCaMP6m signals from nociceptor terminals during heat ramp stimuli. (B and B’) Representative images showing thermal activation of nociceptors in control animals during calcium imaging (*ppk-GAL4 UAS-GCaMP6m* x *w^1118^*). In comparison to the initial frame (B), the GCaMP6m signal monitored at nociceptor axon termini was increased when the probe temperature reached 40°C (B’). (C and C’) Images showing a representative result of animals with nociceptor-specific up-regulation of BMP signaling (*tkv*^Q*D*^ OE, *ppk-GAL4 UAS-GCaMP6m* x *UAS-tkv*^Q*D*^). Compared to the baseline (C), increase of GCaMP fluorescent intensity was observed when the probe temperature reached 40°C (C’). (D) Average percent increase of GCaMP6m fluorescent intensity relative to baseline (ΔF/F_0_) during heat ramp stimulations. ΔF/F_0_ is plotted to binned probe temperature (interval = 1 °C). In controls, GCaMP6m fluorescence in nociceptors began increasing when the probe temperature reached 37 °C, peaked at 43°C, and returned to baseline at 47 °C. In comparison to controls, nociceptors of *tkv*^Q*D*^ OE animals showed a highly exaggerated fluorescent increase of GCaMP through 36°C to 50°C. n = 17 and 19 for controls and *tkv*^Q*D*^ OE, respectively. * p < 0.05, ** p < 0.01, *** p < 0.001 (Mann-Whitney's U-test). (E) Basal GCaMP6m signals (F_0_) did not differ significantly between controls and *tkv*^Q*D*^ OE (n = 17 and 19). p > 0.6 (Mann-Whitney's U-test). All error bars represent standard error.

### Acute up-regulation of BMP signaling induces nociceptive hypersensitivity

Chronic up-regulation of BMP signaling in nociceptors caused sensitization of behavioral nociception responses of larvae and an increased Ca^2+^ response of nociceptors to noxious heat, but also expansion of nociceptor terminals. To further separate the physiological and developmental effects of BMP up-regulation in nociceptors, we acutely up-regulated BMP signaling. Using the temperature sensitive repressor of GAL4 activity (GAL80^ts^) (42), we activated expression of *tkv^QD^* in larval nociceptors by shifting *ppk-GAL4 UAS-Chr2::YFP tub-GAL80^ts^* animals to 30°C for 24 hours. We then tested these larvae for sensitized optogenetic nociception. The acute induction of *tkv^QD^* induced hypersensitivity in the optogenetic nocifensive responses and also significantly increased nuclear pMad levels relative to controls (Fig 7A and B). However, no detectable axonal overgrowth was induced by acute *tkv^QD^* expression (Fig 7C and D). Unfortunately, we were not able to investigate the effects of this manipulation on nociception responses with a 39°C thermal stimulus because the prolonged incubation at 30°C interfered with 39°C NEL behavior in both controls and experimental animals (Fig S5). This latter finding indicates that the sensitivity of thermal nociception in *Drosophila* is modulated by the ambient temperature of the environment. Collectively, these data demonstrate that acute activation of BMP signaling in nociceptors is sufficient to sensitize larval nociceptive response in the absence of detectable changes to axonal morphology. Taken together with our Ca^2+^ imaging results, these data suggest a physiological, rather than a developmental, role for BMP signaling in the regulation of nociceptor sensitivity.

**Fig 7.**
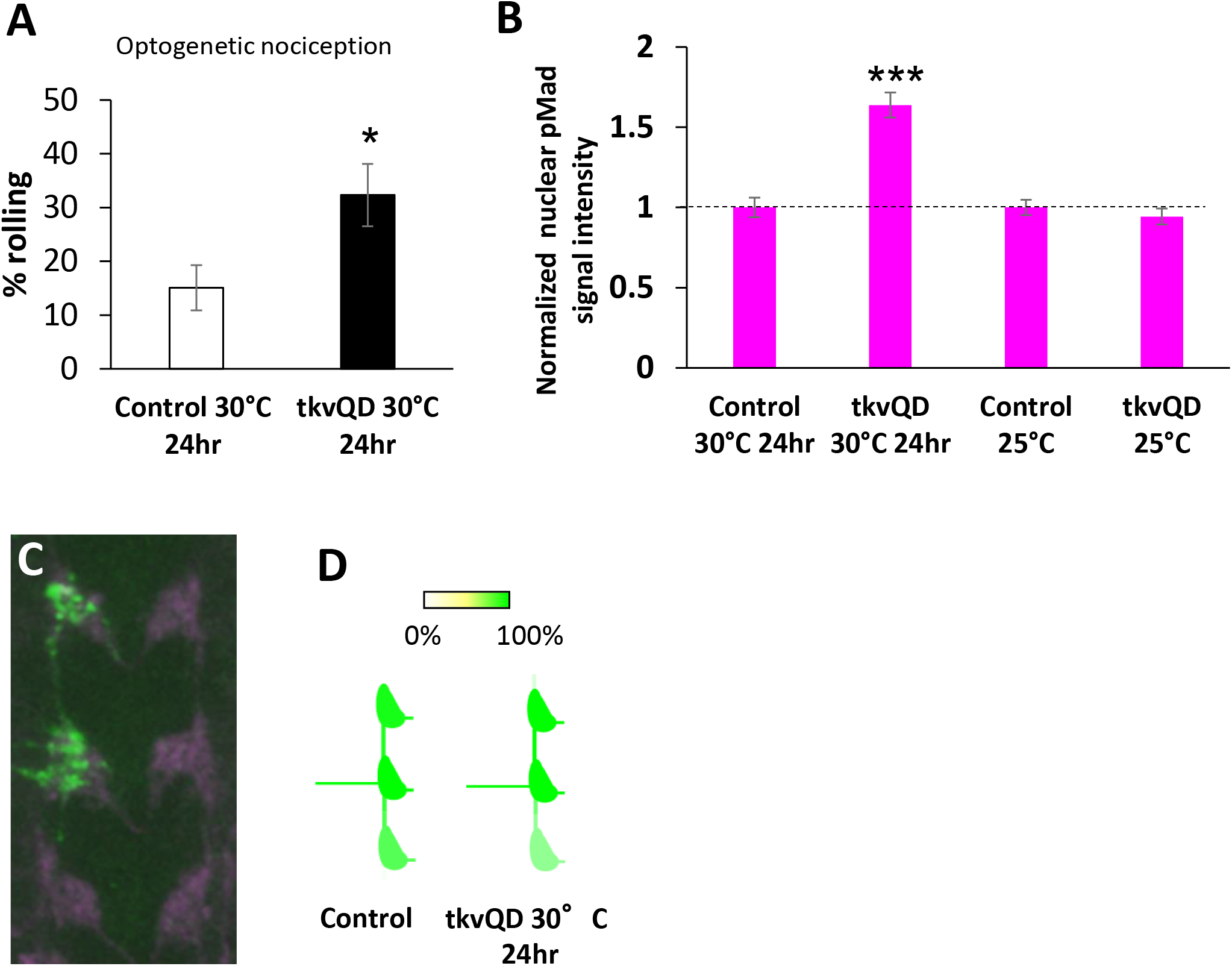
Acute up-regulation of BMP signaling sensitizes optogenetic nociception. (A) Acute expression of *tkv*^QD^ induced hypersensitivity in optogenetic nociception. After 24 hour induction of *tkv*^QD^ in Class IV nociceptors (ppk-GAL4 UAS-ChR2::YFP/+; UAS-*tkv*^QD^/tub-GAL80^ts^ incubated at 30°C for 24 hours, n = 65, 32 ± 6%), larval nociceptive responses to optogenetic activation of Class IV nociceptors were significantly increased compared to those in controls (ppk-GAL4 UAS-ChR2::YFP/+; tub-GAL80^ts^/+ incubated at 30°C for 24 hours, n = 73, 15 ± 4%). * p < 0.05 (Fisher's exact test). (B) Acute induction of *tkv*^QD^ increased nuclear pMad levels in nociceptors. pMad levels in nociceptor nuclei were significantly elevated (64 ± 8%) in animals with 24 hour *tkv*^QD^ induction (ppk-GAL4 UAS-mCD8::GFP/+; tub-GAL80^ts^/UAS-*tkv*^QD^ incubated at 30°C for 24 hours, n = 12) compared to control animals (ppk-GAL4 UAS-mCD8::GFP/+; tub-GAL80^ts^/+ incubated at 30°C for 24 hours, n = 12). When raised at 25°C, animals with UAS-*tkv*^QD^ (ppk-GAL4 UAS-mCD8::GFP/ppk-CD4-tdGFP; tub-GAL80^ts^/UAS-*tkv*^QD^, n = 48) and controls (ppk-GAL4 UAS-mCD8::GFP/ppk-CD4-tdGFP; tub-GAL80^ts^/+, n = 48) showed comparable pMad levels. *** p < 0.001 (Mann-Whitney's U-test). (C and D) 24-hour induction of *tkv*^QD^ did not induce axonal overgrowth. (C) A representative image of axon termini of a single v’ada neuron. Scale bar represents 5 μm. (D) Heat map of v’ada axonal projection. 24-hour expression of *tkv*^QD^ did not cause a severe overextension phenotype (n = 7). The heat map of the control is reused from Fig. 5G for comparison. All error bars represent standard error.

## Discussion

Identifying novel conserved molecular pathways that regulate nociception in model animals is a promising strategy for understanding the molecular basis of pain signaling and pain pathogenesis (43, 44). Using *Drosophila*, we found that both the E3 ligase Hiw and the downstream BMP signaling pathway play crucial roles in regulating nociceptor sensitivity.

### Hiw's complexed roles in regulating nociceptor functions

The data we present in this study suggest that *hiw* has at least two distinct functions in the regulation of nociceptor sensitivity. We found that strong loss-of-function mutants of *hiw* showed insensitivity to noxious heat but hypersensitivity to optogenetic stimulation of nociceptors (Fig 1A and C). Since expressing wild-type *hiw* in nociceptors of *hiw* mutants rescued both phenotypes, loss of *hiw* in nociceptors is responsible for these two ostensibly opposing phenotypes (Fig 1A and C). We also found that nociceptor-specific expression of hiwRNAi or *hiwΔRING* caused only hypersensitivity (Fig 1D and E) (16), indicating that the process that governs hypersensitivity is separable from the cause of insensitivity. As insensitivity was epistatic to hypersensitivity in thermal nociception assays, we used optogenetics to show that hypersensitivity is actually present in *hiw* genetic mutants as well as in previously described RNAi animals. The use of optogenetic stimulation of neurons allowed us to bypass the endogenous sensory transduction step(s) and to reveal this role. Our data suggest that *hiw* is a) required for the negative regulation of a neural pathway that is downstream sensory transduction and b) required to confer normal sensitivity to noxious heat via sensory transduction pathways. As strong *hiw* loss of function causes reduced dendritic arbors (40) while *hiw* RNAi does not (16), it is possible that the reduced dendrite phenotype accounts for the insensitivity of the strong *hiw* alleles. Consistent with this hypothesis, many mutants that cause insensitive thermal nociception are associated with a reduction in the dendritic arbor (16). The phenotypic difference between strong loss-of-function mutants and RNAi or Hiw dominant-negative animals suggests that insensitive and hypersensitive phenotypes observed in *hiw* mutants have different sensitivity to the dosage of *hiw*. This has also been seen in the larval motor neuron system where it has been demonstrated that two different phenotypes of *hiw* in larval NMJ (overgrowth of synaptic boutons and diminished synaptic function) are separable by their different sensitivity to the dosage of *hiw* (20).

Our data also suggest that *hiw* may regulate distinct molecular pathways in motor neurons and in nociceptors. In the larval NMJ, mutations of *hiw* or expression of *hiwΔRING* cause a diminished evoked excitatory junction potential (EJP) due to decreased quantal content in synaptic vesicles (18, 20, 45). However, this diminished evoked EJP phenotype is apparently opposite to the hypersensitive nociception phenotype observed in this study. Thus the downstream targets and/or pathways of Hiw in nociceptors may be distinct from those in motor neurons.

We identified the BMP signaling pathway as an important signaling pathway in nociceptors that is regulated downstream of *hiw*. In fly motor neurons, it has been proposed that BMP signaling is a direct target of Hiw ligase (23). However, a later study reported that pMad up-regulation was not detected in motor neuron nuclei in *hiw* mutants (30) and controversy has arisen over this interaction. We found that nuclear pMad signals were up-regulated in *hiw* mutant nociceptors, and that this molecular phenotype was rescued by wild-type *hiw* expression (Fig 2). In addition, we also detected striking accumulation of pMad in both the nuclei and cytoplasm of nociceptors expressing Hiw dominant negative proteins (Fig 2 and 4). Finally, using *UAS-mad^1^*, we showed that a Mad-dependent pathway is responsible for the hypersensitive thermal nociception caused by *hiwΔRING* expression (Fig 2H). Our data therefore support the idea that the nociceptor BMP signaling pathway is regulated downstream from *hiw*.

Although we demonstrated that BMP signaling is downstream of *hiw* in nociceptors, we have yet to determine the precise mechanism for Hiw regulation of BMP signaling. Our genetic analysis suggests that BMP signaling in nociceptors is regulated independently from the *wnd* pathway (Fig 3). Wnd is the best characterized target substrate of Hiw in the regulation of NMJ morphology (30, 37-40, 46). Our expression analysis using various *hiw* deletion series showed that the set of *hiw* deletion constructs that induced up-regulation of BMP signaling in nociceptors was not identical to the set that induced abnormal synaptic morphology in motoneurons (38). This finding is somewhat consistent with the existence of a Wnd-independent mechanism in the regulation of BMP signaling in nociceptors, since the Hiw-Wnd pathway plays a pivotal role in regulating synaptic morphology in larval NMJ.

Intriguingly, our expression study of the *hiw* deletion series showed that the expression of HiwNT caused a prominent accumulation of nuclear pMad, while the expression of HiwCT or HiwΔRCC1 caused accumulation of pMad signals in both the nuclei and cytoplasm in nociceptors (Fig 4C-E). These data raise the possibility that Hiw is involved in at least two different mechanisms which regulate pMad: one pathway affecting nuclear pMad and another for cytoplasmic pMad. Given that *hiw* is a large protein with many functional domains for interacting with multiple molecules, the notion that *hiw* is involved in multiple processes regulating various aspects of neuronal functions in both motor neurons and nociceptive sensory neurons is perhaps unsurprising. Further studies are necessary to reveal the mechanisms of Hiw-dependent regulation of BMP signaling in nociceptors.

### Physiological effects of BMP signaling in nociceptor axon terminals

We have presented a new physiological preparation for investigating the calcium levels in nociceptor terminals with a physiologically relevant noxious heat stimulus. This allowed us to demonstrate that up-regulation of BMP signaling in nociceptors sensitizes the physiological responses of nociceptors in response to noxious heat in addition to its effects on behavior (Fig 5 and 6). We also demonstrated that acute activation of intracellular BMP signaling in nociceptors is sufficient for the nociceptive sensitization (Fig 7). Although it has been previously reported that BMP signaling in nociceptors is required for nociceptive sensitization after tissue-injury in *Drosophila* (22), the mechanisms of the regulation of nociception by BMP signaling was totally unknown. Our study provides the first evidence that BMP signaling regulates calcium signaling in axon termini of nociceptors.

The BMP signaling pathway plays crucial roles in various developmental processes, such as embryonic patterning, skeletal development, and the development of neuronal circuits (47, 48). The roles of BMP signaling in the regulation of neuronal activity has also been extensively investigated in larval motor neurons, where BMP signaling plays crucial roles in the homeostatic regulation of synaptic morphology and transmission (49, 50). Since BMP signaling is important for synaptic transmission in larval NMJ, an interesting hypothesis is that BMP signaling in nociceptors functions to homeostatically regulate synaptic function (similar to that seen in motor neurons). Again, our study suggests that BMP signaling acts differently in nociceptors than in motor neurons. First, although our data show that activated intracellular BMP signaling in nociceptors resulted in hypersensitivity of nociception and nociceptor sensitivity, genetic manipulations that increase intracellular BMP signaling in motor neurons does not increase evoked EJP in larval NMJ (24, 27-29). Second, interfering with BMP signaling with dominant negative Mad did not cause nociception insensitive phenotypes (Fig S3) (consistent with another study that found that nociceptor-specific knockdown of BMP signaling components did not affect basal thermal nociception (22)). In contrast, loss of BMP signaling components in motor neurons decreased evoked EJP by disturbing homeostatic regulation of synapses (23, 35, 51). Finally, local pMad signals were detected at NMJ as well as at nuclei in motor neurons. This is relevant in that local pMad at the NMJ functions through a non-canonical signaling pathway to regulate synaptic maturation (25, 34). In the case of nociceptors, however, pMad signals were undetectable at nociceptor axon termini (Fig 2G). Although a full understanding of the mechanisms through which BMP signaling regulates nociceptor sensitivity requires further investigation, these results indicate that BMP signaling may act differently in the nociceptors and motor neurons to regulate neuronal outputs.

### Potential conservation of Hiw-BMP pathway in regulating nociception in mammals

Hiw and BMP signaling pathway components are all evolutionally well-conserved. The role of *hiw* in the negative regulation of nociceptive signaling may be as well. A mammalian *hiw* orthologue *Phr1/MYCBP2* has been previously implicated in a negative regulation of nociception processing. Specifically, it has been reported that *Phr1/MYCBP2* is expressed in DRG neurons, and that intrathecal injection of an antisense oligonucleotide against *Phr1/MYCBP2* causes hypersensitivity in formalin-induced nociceptive responses (52). Furthermore, nociceptive and thermoceptive neuron-specific *Phr1/MYCBP2* knock-out mice show prolonged formalin-triggered sensitization in thermal nociception, whereas no obvious phenotypes are observed for basal nociception in the knock-out animals (53). Decreased internalization of the TRPV1 channel (which is mediated through a p38 MAPK pathway) has been implicated in this prolonged nociceptive sensitization in *MYCBP2* knock-out mice (53). In contrast, whether BMP signaling plays a role in regulating nociception in mammals is unknown. Similarly, the degree to which the role of Hiw and BMP signaling is conserved in the physiological regulation of mammalian nociceptors represents a fascinating topic for future investigation.

Intriguingly, Hiw and BMP signaling have been implicated in nerve regeneration/degeneration processes after axonal injury in both *Drosophila* and mammals (17, 54). In flies, axonal injury leads to decrease of Hiw, which leads to the upregulation of Wnd that promotes axonal degeneration in motor neurons (46). Phr1/MYCBP2 is also involved in promoting axonal degeneration after sciatic or optic nerve axotomy (55). Smad1 is known to be activated and play an important role for axonal regeneration after peripheral axotomy of DRG neurons (56-59). Because nerve injuries are thought to be one of key contributors for neuropathic pain conditions and peripheral axotomies are widely used to generate neuropathic pain models in mammals, it will be of particular interest in the future to determine whether the Hiw-BMP signaling pathway and up-regulation of intracellular BMP signaling in nociceptors play a role in the development of a neuropathic pain state in mammals.

## Materials and Methods

### Fly strains

Canton-S and *w^1118^* were used as control strains as indicated. The other strains used in this study were as follows: *ppk1.9-GAL4* (60), UAS-*mCD8::GFP* (61), UAS-*ChR2::YFP* line C (4), *hiw*^*ND8*(18)^, *hiw^ΔN^*, *hiw^ΔC^*, UAS-*hiw*, UAS-*hiwΔRing* (20), UAS-*hiwNT, UAS-hiwCT*, UAS-*hiwΔRCC*, UAS-*hiwΔHindIII*, UAS-*hiwCT1000* (38), *wnd*^1^, *wnd*^2^, UAS-*wnd* (30), *ppk1.9-GAL4; UAS>CD2 stop>mCD8::GFP hs-flp, UAS-tkv*^QD^ (33), *tub-GAL80^ts^* (62), *ppk-CD4-tdGFP* (63) and UAS-*GCaMP6m* (41). UAS-*mad^1^* (36)

### Thermal nociception assay

The thermal nociception assay was performed as described previously (3, 6, 10, 16, 64). NEL latency was measured as initial contact of the thermal probe on the lateral side of the larval body wall to the completion of NEL (a 360° roll). Stimulation was ceased at 11 seconds. A thermal probe heated to 46°C was used to examine the insensitive phenotype since it usually evokes NEL in less than 3 seconds (3, 6, 10, 16, 65). A 39°C probe, which usually results in NEL in 9-10 seconds, was used to examine thermal hypersensitivity, as using a lower temperature probe is important to detecting the hypersensitive phenotype (16).

### Optogenetic nociception assay

The optogenetic nociception assay was performed as described previously (5) with slight modifications. 3.8 klux was used to test for optogenetic hypersensitivity, but 76 klux blue light was used in the analysis of acute *tkv*^QD^ induction (Fig 7). Because male larvae show a lower responsiveness to optogenetic nociceptor activation than females (Honjo, unpublished), male larvae were used to allow for more easily detectable hypersensitivity.

### Immunohistochemistry

Antibodies used in this study were as follows: rabbit anti-GFP (Invitrogen, 1:1000), mouse anti-GFP (Invitrogen, 1:250), mouse anti-rat CD2 (AbD Serotec, 1:200), rabbit anti-pMad (gift from Ed Laufer, 1:1000), goat anti-rabbit Alexa488 (Invitrogen, 1:1000), goat anti-rabbit Alexa568 (Invitrogen, 1:1000), goat anti-mouse Alexa488 (Invitrogen, 1:1000) and goat anti-mouse Alexa568 (Invitrogen, 1:1000). Larvae were filleted, fixed in 4% paraformaldehyde for 30 minutes and then stained according to standard protocols.

### pMad staining and image analysis

Wandering third instar larvae expressing mCD8::GFP in nociceptors were filleted and immunostained as described above. To minimize variation due to processing controls, experimental specimens were processed side-by-side within the same staining solutions. The dorsal Class IV mutidendritic neuron (ddaC) was imaged in segments A4-6 (Zeiss LSM 710 with a 100x/1.4 Plan-Apochromat oil immersion or Olympus FV1200 with a 100x/1.4 UPLSAPO oil immersion). Z-stack images were converted to maximum intensity projections. To quantify nuclear pMad signals, nociceptor nuclei were identified based on the absence of GFP signal, and a region of interest (ROI) outlining the nucleus was delineated. The average signal intensity of nuclear pMad staining in the ROI was then calculated. Background signal intensity was determined as the mean from ROIs (identical size and shape of the nucleus from the image) drawn in the four corners of each image. The calculated background signal intensity was then subtracted from the nuclear pMad signal level. Data are plotted as nuclear pMad levels normalized to that of the co-processed control specimens. Image analyses were performed in Adobe Photoshop.

### Dendrite imaging and quantification

Wandering third instar larvae expressing mCD8::GFP in nociceptors under the control of ppk1.9-GAL4 were anesthetized by submersion in a drop of glycerol in a chamber that contained a cotton ball soaked by a few drops of ether. ddaC neurons in segments A4-6 were imaged on Zeiss LSM 5 Live with a 40x/1.3 Plan-Neofluar oil immersion objective lens. A series of tiled images were captured and assembled to reconstruct the entire dendritic field of the three A4-6 ddaC neurons. Z-stack images were then converted to maximum intensity projections. Dendritic field coverage was quantified as described previously (16).

### Flip-out clone analysis of axon termini

A *ppk1.9-GAL4; UAS>CD2 stop>mCD8::GFP hs-flp* strain was used to induce single cell flip-out clones in order to sparsely label nociceptors. Six virgin females and three males were used to seed vials containing a cornmeal molasses medium for a period of 2 days. The seeded vials were then heat-shocked in a 35°C water bath for 30 minutes. After an additional 3 to 5 days, wandering third instar larvae were harvested from the vials and dissected. In order to precisely identify the neurons responsible for the axons labeled in the CNS, the incision made in filleting the larvae was along the dorsal side, and the CNS remained attached to the fillet prep during immunostaining. mCD8::GFP and rat CD2 were detected using rabbit anti-GFP and mouse anti-rat CD2 primary antibodies, and visualized by anti-rabbit Alexa488 and anti-mouse Alexa568 secondary antibodies, respectively. Axon terminal branches of single cell flip-out clones were imaged in the abdominal ganglion using a Zeiss LSM 5 Live with a 40x/1.3 Plan-Neofluar oil immersion objective. The cell body of origin for each flip-out clone was then determined by inspecting the body wall of the corresponding fillet. Flip-out clones belonging to A1-7 segments were imaged and analyzed.

To analyze the projection patterns for axon terminals, the presence or absence of terminal branches in each neuromere and longitudinal tract was manually identified for each single nociceptor clone. In order to align clones projecting to different segments, positions relative to the entry neuromere were used. The neurons that aligned were then used to calculate the percentage projecting to each neuromere and longitudinal tract. Heat-maps were color-coded according to these percentages using Microsoft Excel and Adobe Illustrator.

The quantification of axon terminal area was performed in Matlab. Z-stack images of axon termini were converted to maximum intensity projections and manually cropped to exclude signals from other clones in the same sample. The green channel (GFP) and red channel (CD2) of the cropped images were separately binarized using Otsu's method (66). The number of GFP-positive pixels were counted to calculate the area innervating the termini. To compensate for differences in the size and shape of the ventral nerve cord, the number of GFP-positive pixels was normalized to the average size of a single neuromere, which was calculated as the number of CD2-positive pixels divided by the number of neuromeres in the cropped image.

### Acute induction of *tkv*^QD^ by tub-GAL80^ts^

Larvae raised in normal fly vials for 5 or 6 days at 25°C, or larvae raised on apple juice plates containing ATR for 4 days at 25°C, were transferred to 30°C for 24 hours. In every experiment, experimental genotypes and control animals were treated side-by-side to minimize the effect of potential variations in temperature.

### Calcium imaging

*The ppk1.9-GAL4 UAS-GCaMP6m* strain was crossed to either a control strain (*w^1118^*) or UAS-*tkv^QD^* strain. Activity of larval nociceptors were monitored at their axon terminals in the larval ventral nerve cord (VNC), which was exposed for imaging by a partial dissection as follows: wandering third instar larvae expressing GCaMP6m in Class IV md neurons were immobilized in ice cold hemolymph-like saline 3.1 (HL3.1) (70 mM NaCl, 5mM KCl, 1.5 mM CaCl_2_, 4 mM MgCl_2_, 10 mM NaHCO_3_, 5 mM Trehalose, 115 mM Sucrose, and 5 mM HEPES, pH 7.2)(67). The outer cuticle of each larvae was cut at segment A2/A3 to expose the central nervous system from which intact ventral nerves innervate the posterior larval body. The partially dissected animals were transferred to an imaging chamber containing HL3.1 equilibrated to the room temperature (23-25 °C). A strip of parafilm was placed over the larval VNC and was used to gently press the nerve cord down onto a coverslip for imaging. A Zeiss LSM5 Live confocal microscope and a 20x/0.8 Plan-Apochromat objective with a piezo focus drive were used to perform three-dimensional time-lapse imaging. Z-stacks consisting of 10-11 optical slices (Z depth of 63 to 70 μm) of 256 x 128 pixel images were acquired at approximately 4 Hz. During imaging, and using a custom-made thermal probe, a heat ramp stimulus was applied locally to one side of the A5 to A7 segments. The temperature of the thermal probe was regulated using a variac transformer. 10V was used to generate a 0.1 °C/sec heat ramp stimulation and no voltage was applied during cooling. A thermocouple probe (T-type) wire was placed inside of the thermal probe to monitor the probe temperature, and the data were acquired at 4 Hz through a digitizer USB-TC01 (National Instruments) and NI Signal Express software (National Instruments). The acquired images and temperature data were analyzed using Matlab software (Mathworks). Maximum intensity projections were generated from the time-series Z-stacks. Region of interest (ROI) was selected as a circular area with a diameter of 6 pixels, whose center was defined as the centroid of the A6 neuromere. Averaged fluorescent intensities (F) were calculated for the ROI for each time point. The average of Fs from the first 30 frames was used as a baseline (F_0_), and the percent change in fluorescent intensity from baseline (ΔF/F_0_) was calculated for each time point. Since acquisitions of images and probe temperatures were not synchronized, probe temperature for each time point was estimated by a linear interpolation from the raw probe temperature reading. For a comparison of controls and *tkv^QD^* OE, ΔF/F_0_, data were binned and averaged in 1°C intervals.

### Statistical analyses

To statistically compare proportional data, Fisher's exact test was used. Multiple comparisons of proportional data were corrected by the Bonferroni method. For non-proportional data, Mann-Whitney's U-test was used for pair-wise comparisons, and Steel's test (non-parametric equivalent of Dunnet's test) was used for multiple comparisons. Statistical analyses were performed in R software and Kyplot.

## Acknowledgements

We thank TRiP at Harvard Medical School (NIH/NIGMS R01-GM084947), Aaron DiAntonio, Michael B. O’Connor, Gary Struhl, Chunlai Wu, and the Bloomington Stock Center for fly stocks. We are grateful to Dan Vasiliauskas, Susan Morton, Tom Jessell, and Ed Laufer for kindly providing the pMad antibody. We also thank Dr. Emiko Suzuki for her mentorship and helpful advice to KH. We acknowledge Hisako Honjo for technical assistance in dendrite analysis. Stephania Mauthner, Andrew Bellemer, and Melissa Christiansen made helpful suggestions on this manuscript. This work was supported by a grant from the National Institutes of Health (R01GM086458, W.D.T.) and JSPS KAKENHI (26890025, K.H.). K.H. was supported by postdoctoral fellowships from the Uehara Memorial Foundation, the Ruth. K. Broad Biomedical Research Foundation, the Japan Society for the Promotion of Science, and the National Institute of Genetics.

